# Spatial ecology and population dynamics of brown trout *Salmo trutta* L. in reservoirs and headwater tributaries

**DOI:** 10.1101/2023.12.06.570412

**Authors:** Jamie R. Dodd, Richard A.A. Noble, Andy D. Nunn, Holly Owen, Paolo Moccetti, Jonathan P. Harvey, Liam Wallace, Ben Gillespie, Domino A. Joyce, Jonathan D. Bolland

**Author notes:** Author for correspondence:School of Natural Sciences, University of Hull Cottingham Road, Hull, HU6 7RX, UK. This investigation was funded by Yorkshire Water Services Limited.

## Abstract

This investigation compared the spatial ecology and population dynamics of brown trout *Salmo trutta L*. between reservoirs with (impact; Langsett Reservoir) and without (control; Grimwith Reservoir) barriers to fish movements into afferent headwater tributaries, including the effectiveness of a fish pass to remediate connectivity. Passive Integrated Transponder (PIT) telemetry revealed fish that emigrated from Langsett and Grimwith reservoirs were 1-3 and 0-2 years old, respectively, and predominantly did so in March-May and October-December in both. Weirs at Langsett Reservoir (emigration rate = 26%) appeared to thwart emigration relative to Grimwith Reservoir (emigration rate = 85%). Acoustic telemetry (2D positions) in the impact reservoir revealed the largest home range was in October – December (monthly K95 ± S.D. up to 26.9 ± 6.69 ha in November), activity was influenced by both month and time of day, and fish occupied shallow water depths (relative to reservoir depth), especially at night. Large proportions of brown trout tagged in Grimwith and Langsett reservoirs (42.9% and 64.1%, respectively) and fish that emigrated (37.2% and 27.7%, respectively) were detected moving upstream into tributaries. At both reservoirs, peak immigration for 3- and ≥4-year-old fish occurred in October-December, although upstream movements occurred throughout the year and by all age classes. Three brown trout passed upstream of each of the weirs on River Little Don (prior to fish pass construction; 3% of those that approached from downstream) and Thickwoods Brook (throughout the study; 2%). Overall fish pass solution passage efficiency was 14% but was higher for 2- and 3-year-old fish (32%), which was comparable to fish translocated from upstream (33%). Passage predominantly occurred at lower river levels than fish pass entrance / attraction, which was also lower than during approaches to the weir. A Before-After Control-Impact (BACI) design found that although juvenile (0+, but not >0+) brown trout densities were lower after fish pass construction, the reduction was significantly less than at control sites, i.e., the fish pass had a positive effect. Overall, this investigation significantly furthers our understanding of brown trout spatial ecology and population dynamics in reservoirs and headwater tributaries.

## 1. INTRODUCTION

Brown trout *Salmo trutta* L. is one of the most widespread and extensively studied fish species. It is native to Europe, western Asia and North Africa, but has also been widely introduced elsewhere (McIntosh *et al*., 2012). A key aspect of the species’ ecology is the existence of two contrasting life history strategies, namely freshwater residency and anadromy (Klemetsen *et al*., 2003). In the resident form (brown trout), the entire life cycle is completed in freshwater, whereas anadromous individuals (sea trout) migrate between marine and freshwater ecosystems. Irrespective of the life history strategy, individuals invariably need to migrate between habitats according to temporal or ontogenetic requirements. For anadromous individuals, this necessarily involves movements between marine and freshwater environments, whereas freshwater residents migrate within rivers or between rivers and still waters (Arostegui & Quinn, 2019; Ferguson *et al*., 2019).

Riverine migrations have been the most extensively studied, with upstream movements to spawning grounds frequently triggered by elevated flows in autumn and winter (Klemetsen *et al*., 2003; Jonsson & Jonsson, 2011). By contrast, a combination of competition and genetic drivers instigates downstream migrations to habitats offering superior feeding opportunities, such as larger rivers, still waters and the sea (Arnekleiv *et al*., 2007; Jonsson & Jonsson, 2011). Compared to rivers, much less is known about the migrations of brown trout that inhabit still waters, and especially artificial still waters, such as reservoirs which landlock brown trout populations. This is important because a large number of watercourses across the species’ range have been dammed for hydropower, flood prevention, recreation and water supply, thus creating reservoirs (Yasarer & Sturm, 2016). Although brown trout can inhabit and, indeed, flourish in reservoirs, access to tributaries is required for successful spawning to occur in the majority of cases (Crisp *et al*., 1984). When migrations between the reservoir and afferent headwater tributaries are impeded by man-made structures the impacts are potentially greater than in rivers, with monodirectional movements over the weir causing fish populations upstream to become impoverished and the reservoir acting as a population sink.

One option to mitigate the barrier effects of man-made structures (Humphries & Winemiller, 2009; Liermann *et al*., 2012; Dias *et al*., 2017) is construction of fish passes. The performance of fish passes is highly variable and studies on brown trout have produced mixed results (Forty *et al*., 2016; Dodd *et al.,* 2017, 2018, 2023; Mameri *et al.,* 2019; Lothian *et al*., 2020). Even for a single species, passage efficiency is influenced by a range of factors, including fish pass design (e.g., water velocity and turbulence), individual motivation, body length (swimming ability), water temperature and opportunity (Noonan *et al*., 2012; Albayrak et al., 2020; Cano-Barbacil et al., 2020). In addition, the wide range in water levels experienced in reservoirs can have a significant influence on the ability of fish to access tributaries, and there is also potential for differences in the physiological fitness (swimming performance) of individuals inhabiting lentic (reservoirs) vs. lotic (tributaries) environments, which could have implications for passage success at migration barriers. Ultimately, facilitating access to headwater tributaries in reservoirs will ensure spawning success and likely lead to a population size to increase, but is rarely quantified.

In contrast to the considerable volume of research that has been conducted on brown trout in rivers, there is a paucity of information on the species in reservoirs and their tributaries. Thus, this study used acoustic telemetry to quantify the spatial ecology and activity of brown trout in a typical upland reservoir in the species’ native range. Passive integrated transponder (PIT) telemetry was used to examine longitudinal movements in afferent headwater tributaries fragmented by man-made weirs (i.e., impact), focusing on the influence of fish age, timing of movements and tagging location. These results were interpreted in the context of a concurrent study at a reservoir where brown trout had unimpeded access to headwater tributaries (i.e., control). In addition, the efficiency of a fish pass constructed mid-study was assessed, focusing on the influence of fish age and river level, including fish translocated from upstream of the weir (Dodd *et al*., 2023). The impact of the fish pass on the brown trout population upstream was quantified using a BACI (Before-After, Control-Impact) assessment, to increase the ability to account for and differentiate treatment effects from natural temporal variability (Roni & Beechie, 2013; Angelopoulos et al., 2017; Mahlum et al., 2018). It was hypothesised that the fish pass would remediate the immigration of large, and therefore more fecund (Klemetsen *et al*., 2003), individuals from the reservoir into the tributary, with a consequent increase in the brown trout population over time.

## 2. MATERIALS AND METHODS

### 2.1 Study area

Langsett Reservoir (53.4968: -1.6860; 50.5 ha) and Grimwith Reservoir (54.0771, -1.9094; 147 ha) are situated in northern England and used for drinking water supply (Yorkshire Water) (Figure 1). Both were classified as heavily modified water bodies having a “moderate” ecological potential under the WFD (UKTAG, 2008; EA, 2019). Langsett Reservoir is primarily fed by two headwater tributaries, the River Little Don (RLD) and Thickwoods Brook (TWB), both of which have large siltation weirs at their confluence with the reservoir (Figure 1). These weirs reduce the longitudinal connectivity of the system, thus isolating the reservoir’s fish populations from those in the tributaries and having possible impacts on their ecological status. Grimwith Reservoir is primarily fed by three tributaries, Blea Gill Beck (BGB), Gate Up Gill (GUG) and Grimwith Beck (GB); both Blea Gill Beck and Gate Up Gill had open access to the reservoir, whilst Grimwith Beck was fragmented by a siltation weir.

**Figure 1.**
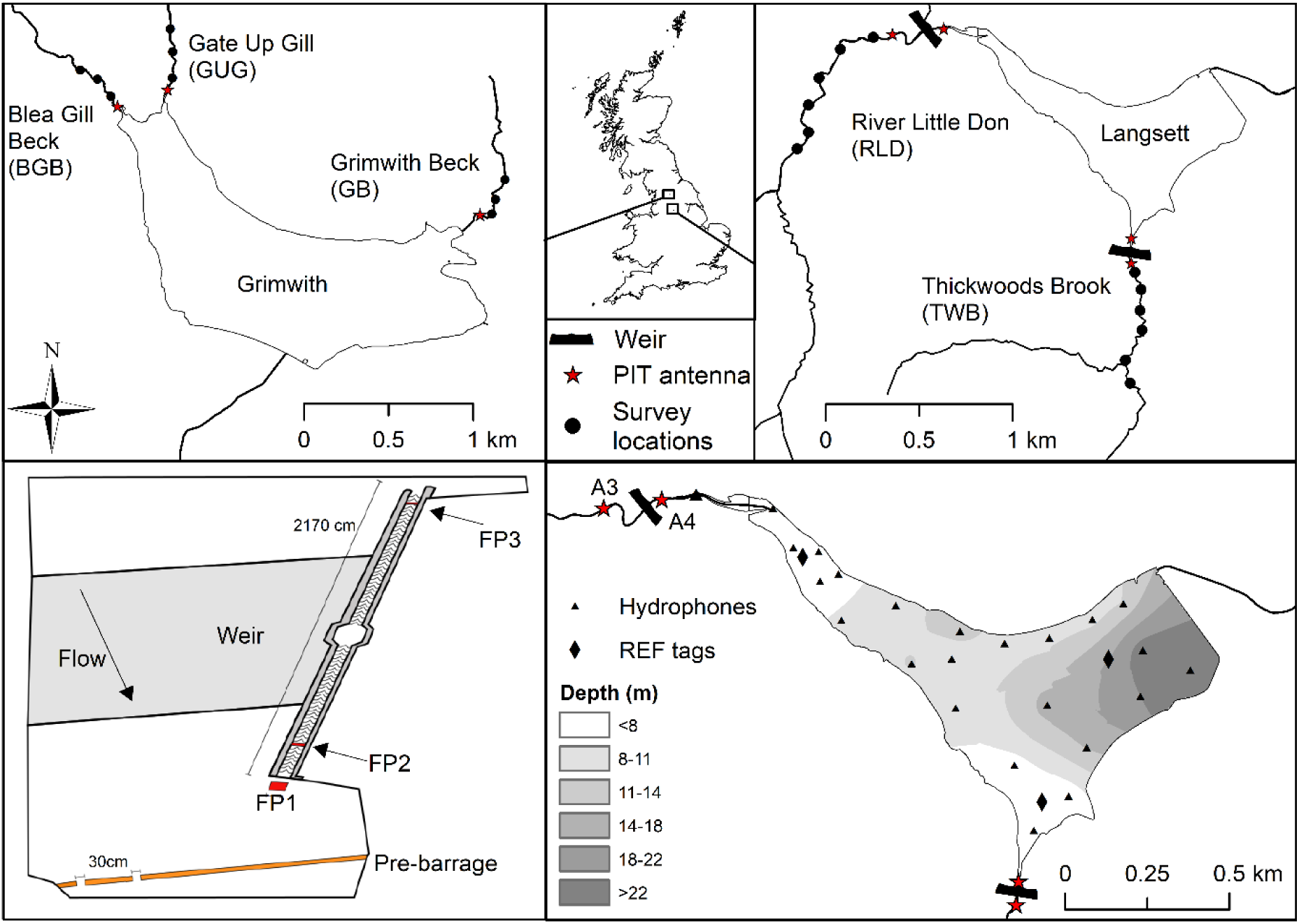
Location of Grimwith and Langsett reservoirs and tributaries with positions of weirs (black rectangles), fish population surveys (black dot), passive integrated transponder (PIT) antennas (red star/boxes), hydrophones (triangles) and reference tags (diamonds).

In December 2017, a two-flight Larinier-style fish pass was opened on the River Little Don weir at Langsett Reservoir (Figure 1). The entrance to the fish pass was located on the left bank, 5 m downstream of the weir face, and a ∼25-cm high low-cost baffle pre-barrage (LCB), used to retain water between the baffle and weir face, was located a further 6 m downstream, creating a pool around the fish pass entrance. The LCB had two 30-cm notches situated close to the right bank to allow for passage past the pre-barrage. The fish pass was operated from 1 October-31 April, the main brown trout migration period, for the remainder of the study, and closed from May-September to maintain the flow of water over the face and, hence, the aesthetic qualities of the weir.

### 2.2 Sampling strategy and data collection

Quantitative three-catch depletion electric fishing surveys were carried out at a total of 21 sites on tributaries upstream of Langsett and Grimwith reservoirs every September/October from 2014–2021. Site length (m) and width (m) were measured to calculate the survey area (m^2^). In addition, brown trout were captured from the reservoirs using seine nets (150 m x 4 m, 20 mm mesh; 2014-2019), fyke nets (2.75 m x 0.53 m, 6 m leader, 10, 14 & 17 mm mesh; 2014-2018) and electric fishing (2015-2021). All fish were measured (fork length, mm) and a scale sample was collected for age determination.

### 2.3 Immigration and emigration

Brown trout were PIT tagged to estimate emigration and immigration rates between the tributaries and reservoirs (up to 101 per tributary per year; 3049 during study). Prior to tagging, fish were checked for any signs of external damage or abnormal behaviour and scanned with a handheld PIT reader to determine if they had been previously tagged. Any fish that had previously been tagged were measured and immediately released, with the tag number and re-capture date and location recorded. All other individuals were anaesthetised using buffered tricaine methanesulphonate (MS-222; 0.8 g/10 L). Once anaesthetised, the fish were measured and a scale sample was taken. Each fish was tagged with a FDX PIT tag (12.0 mm long x 2.1 mm diameter, 0.1 g weight in air, Biomark inc.). The tags were tested with a handheld reader before being injected into the body cavity through a ventro-lateral incision made with a stainless-steel gauge 12 needle, anterior to the muscle bed and pelvic fins. Following tagging, fish were held and monitored in a recovery tank until they had regained balance and began to actively swim. In 2019, a sub-sample of fish (*n* = 26) caught in the River Little Don were displaced downstream of the weir to examine return rate/passage, but all other individuals were released at their capture location. All fish were treated in compliance with the UK Animals in Scientific Procedures Act 1986 under Home Office licence number PPL 60/4400 and PD6C17B56. From 2014-2019, 465 brown trout were PIT tagged and released into Grimwith Reservoir with 1296 released into its tributaries, And between 2014-2021 349 trout were tagged and released into Langsett Reservoir and 939 into its tributaries.

Ten cross-channel pass-over antennae were in tributaries, and two swim-through and one pass-over PIT antennae were at the fish pass. Each station was full-duplex and powered by banks of either 110 Ah or 220 Ah deep-cycle lead-acid batteries connected in parallel. Each station’s batteries were charged by adjacent solar panels. At Grimwith Reservoir, two antennae were located at the mouths of each of the tributaries (when the reservoir was at maximum capacity), enabling the direction of movement of the fish (upstream or downstream) to be recorded. Pairs of antennae in the Grimwith tributaries were ∼15 m apart. At Langsett Reservoir, pass-over antennae were located upstream and downstream of the weir in each tributary. One swim-through antenna was located 0.6 m downstream of the upstream exit of the fish pass and was installed in December 2017, with a second (2.25 m upstream of the fish pass entrance) and a swim-over antenna (0.4 m downstream of the entrance) installed in October 2020 (Figure 1).

The PIT antennae at Grimwith Reservoir were operated from September 2014–February 2020. At Langsett Reservoir, the PIT antennae in the River Little Don and Thickwoods Brook ran from September 2014– March 2022, except for gaps at Thickwoods Brook between November 2018-September 2019 (vandalised and stolen) and February-August 2021 (flooding). Tag detection range at all sites (20-40 cm above the river bed for pass-over antennae; 30 cm either side of the vertical plane for swim-through antennae) was tested during installation and on each visit (approximately once a month) to ensure the read range of the interrogated water column had not decreased. Tag detections on each loop consisted of date, time, detection period, unique tag ID number and loop number, and were manually downloaded from SD cards in the data logger during site visits.

Fish were deemed to have emigrated from a tributary to the reservoir if detected on either the most downstream antenna or on an antenna in a different tributary. Minimum emigration rates were calculated from fish that were detected on the upstream and then downstream antenna as a proportion of those detected only on the upstream antenna (does not include fish that missed the upstream antenna). Immigration was deemed to have occurred when a fish was detected on any antenna in a different tributary to its capture location or the same tributary 30 days after it was last detected on the downstream antenna. Fish that made multiple immigrations during the study were counted more than once, depending upon the metric being analysed. For example, a fish immigrating in both October and December in the same year would be counted twice in an analysis of immigration timing, but only once in an analysis of immigrant age. Fish age classes were calculated from scale readings and ages were assigned to fish based on their length at tagging. Additional year(s) were added onto fish for each 30 April that passed during detection (co-inside with fish pass shutting and time of emergence). Age classes were determined from modal distributions in the length-frequency histograms to set upper and lower length limits per age class.

### 2.4 Spatial ecology and activity in the reservoir

In September and October 2016, nine brown trout from Langsett Reservoir were acoustic (V9AP accelerometer and pressure tag (48 mm long × 9 mm diameter, 6.6 g weight in air, InnovaSea (previously Vemco), 40-60 second ping rate)) and PIT tagged to monitor spatial ecology and activity in the reservoir and tributary confluences. Each acoustic ping would provide data on current depth (m) or general activity (m s^-2^) at the point of transmission. Prior to surgery, acoustic and PIT tags were tested with handheld detectors, sterilised with betadine and rinsed with distilled water, then inserted into the body cavity through a ventro-lateral incision made with a scalpel, anterior to the muscle bed and pelvic fins, and the incision was closed with an absorbable monofilament suture. After surgery, fish were held in an aerated, water-filled container until they had regained balance and were actively swimming, then released back into the reservoir at their capture site. Two of the fish stopped moving 8 days after tagging, so were removed from the analyses. The movements of the remaining seven brown trout were studied until 31 May 2017.

An array of 23 VR2W/VR2Tx-69 kHz acoustic receivers (InnovaSea (previously VEMCO), Halifax, Canada) and three reference tags (with the same ping strength as the fish tags) were deployed at known locations around Langsett Reservoir to provide a comprehensive coverage of the study area, enable triangulation of acoustically tagged brown trout and validate array performance (Figure 1). Sync tags were attached to each receiver to determine performance and correct for ‘clock drift’; when receiver clocks may run at slightly differing times. A depth measurement was taken during the deployment and battery change of each receiver, which was used to calculate an estimated depth profile around the reservoir using the kriging tool in ArcGIS 10.

Data were quality controlled to remove any triangulated pings where the horizontal positioning error (HPE) value was greater than 30 (Roy *et al*. 2014). The mean measured positioning error of the reference tags where HPE was < 30 was 2.01 m. Home ranges were calculated in RStudio, version 4.0.2 (R Core Team, 2020), using the *latticeDensity* package (Barry & McIntyre, 2011) with nodes spaced 10 m apart. The *latticeDensity* package was selected over the standard kernel density estimate often used to calculate home range as it accounts for the irregular boundaries around the reservoir. The package uses a network of interconnected nodes to form a lattice over the area of the reservoir, with trout triangulations interpolated over the top. This method ensures that home range calculations do not include any areas that are inaccessible to the tagged fish. Monthly mean K50 and K95 values were calculated for each fish to compare home ranges during different months.

Depth in the water column and activity data were tested to see if they fitted the assumptions of normal distribution and homogeneity of variances using qqplots, the Shapiro Wilks test and the Bartlett test. This confirmed that the data were non-normal, therefore the non-parametric Scheirer-Ray-Hare test was used to test for significant differences in activity and depths in the water column between months and times of day, and whether they interacted. When comparisons of monthly variations in activity or depth in the water column were undertaken, a minimum of 10 days per month of data per fish was used. The analysis was conducted in RStudio, version 4.1.3 (R Core Team, 2020), using the *Lattice* (Sarkat, 2008) and *Car* (Fox & Weisberg, 2019) packages.

### 2.5 Fish passage performance

To evaluate the efficiency of the fish pass on the River Little Don, a number of metrics were calculated for when the fish pass was open (1 October-30 April):

- “Available fish” was the number of tagged fish detected on the antenna downstream of the weir in the River Little Don (i.e., A4; Figure 1) that were deemed to have approached from downstream, i.e., not including fish moving downstream towards the reservoir.
- “Attraction efficiency” was the number of fish detected on the antenna immediately downstream of the fish pass entrance (FP1) as a proportion of those that were available (A4).
- “Entrance efficiency” was the number of fish detected on the antenna in the lower flight of the fish pass (FP2) as a proportion of those detected at the entrance (FP1).
- “Larinier passage efficiency” was the number of fish detected on the antenna at the upstream end of the fish pass (FP3) as a proportion of those last detected in the lower flight (FP2).
- “Fish Passage Solution (FPS) passage efficiency” was the proportion of available fish (A4) that successfully ascended the weir via the fish pass (FP3); used as there were initially no attraction or entrance antennae.
- “Overall passage efficiency” was the number of fish detected above the weir (FP3/A3) as a proportion of those last detected or tagged downstream of the weir (A4), and thus incorporates fish that may have ascended the weir using a route other than the fish pass.
- “Overall time to pass” was the time difference between approaching the weir during passage (i.e., the last detection on A4) and ascending (i.e., the first detection at the upstream end of the fish pass, FP3).

Air and water pressure data were collected on an hourly interval using Wireless Wildlife probes for the River Little Don in 2020/21. River depth (cm) was calculated by subtracting air pressure from river pressure. Missing data (equipment fault) were modelled (*r*^2^ = 0.89) using a nearby river level monitoring station (River Don, Bower Hill Bridge, Oxspring; 53.514958, - 1.5907915). River depth exceedance values (*Q*_x_) were calculated for when the fish pass was open to assess the approach, attraction, entrance and passage limitations of the fish pass solution.

### 2.6 BACI assessment of brown trout populations in reservoir tributaries

Estimates of 0+ and >0+ brown trout abundance in the reservoir tributaries were derived from the quantitative electric fishing survey data using a three-catch maximum likelihood removal method (Carle & Strub, 1978) and expressed as numbers/100 m^2^. A BACI analysis was then undertaken to look at the effect of the fish pass on fish populations in the River Little Don (impact site), with the remaining tributaries used as controls. The fish pass opened in December 2017, and thus improved recruitment could potentially occur from 2018. Impact assessment calculations identify the mean change in the parameter assessed and whether a significant effect occurred by calculating confidence limits of the change (t-statistic; P < 0.05). Since all >0+ brown trout caught in 2018 would have been recruited prior to the opening of the fish pass, density data from 2018 were not included in the “after” dataset when considering changes in >0+ densities.

A General Linear Model (GLM) was utilised to perform the BACI analysis. This allowed a greater understanding of how the sources of variation contribute to the eventual outcome of the fish pass installation. To account for the large variability in density estimates for both 0+ and >0+ brown trout at impact and control sites, all density data were transformed (natural log + 1). The premise of this model was to determine a significant interaction between the two fixed variables of Area (control and impact) and Period (before and after) to determine significant effect on the geometric mean density. To reduce the levels of residual error, the random effects of Site and Year were nested within the fixed factors of Area and Period.

## 3. RESULTS

### 3.1 Downstream migration; tributary emigration

Of the 545 brown trout that emigrated from tributaries at Grimwith Reservoir, 27%, 49% and 16% were 0, 1 and 2 years old, respectively, at the time of emigration (Figure 2). Where age could be calculated (*n =* 68), 7%, 29%, 16% and 30% of brown trout that emigrated from tributaries at Langsett Reservoir were 0, 1, 2 and 3 years old, respectively. At Grimwith Reservoir, 36% and 35% of emigrations were in March-May and October-December, respectively, in comparison to 21% and 43% at Langsett Reservoir (Figure 2). Twenty-six percent of brown trout detected upstream of a weir at Langsett Reservoir (*n =* 29/112: CI 19-35%: Thickwoods Brook or River Little Don during periods when both antennae were *in situ*) were subsequently detected downstream, in comparison to 85% (*n* = 382/452; 81-88%) at Grimwith Reservoir.

**Figure 2.**
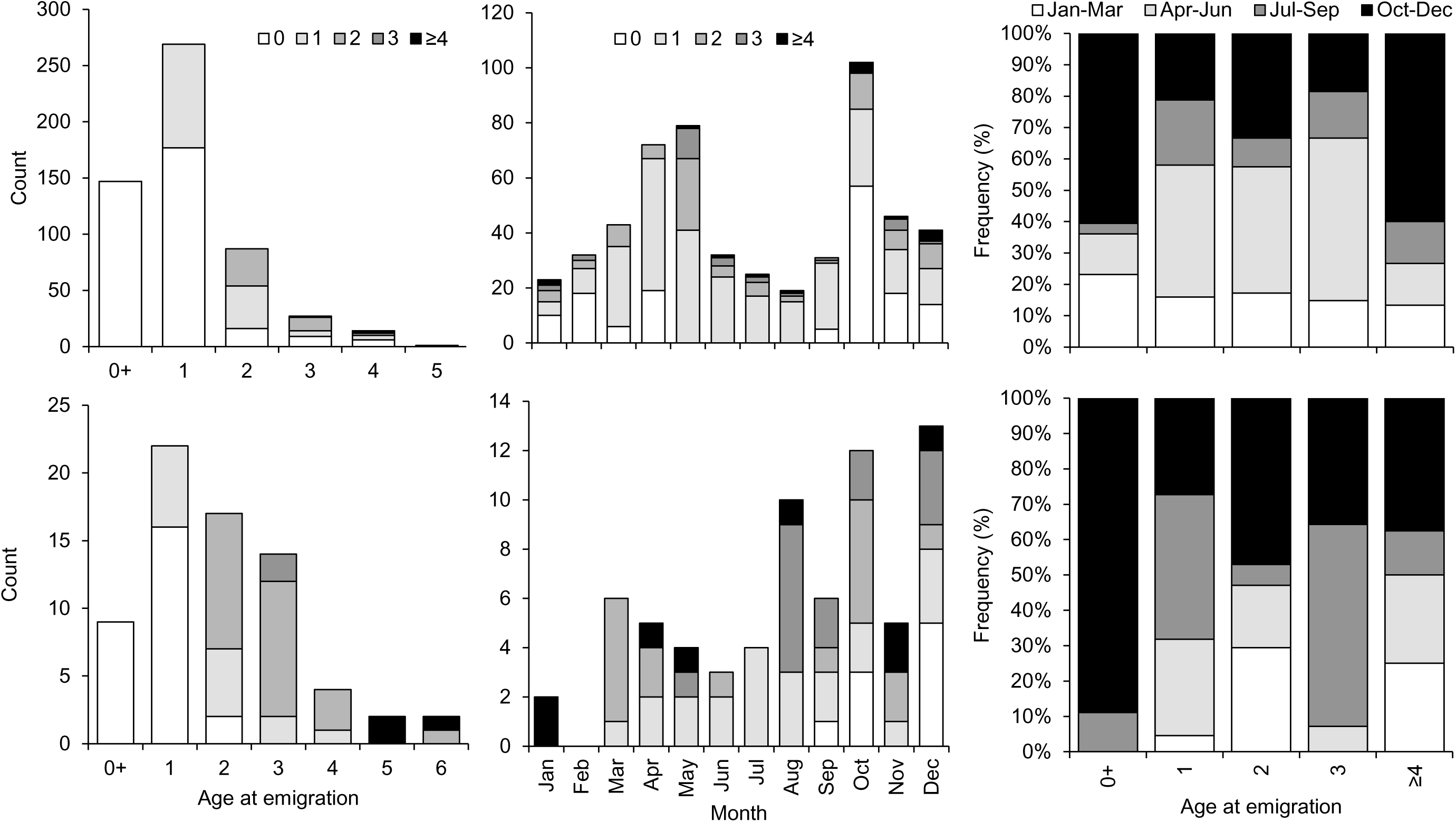
The age (including age at tagging; left) and month (including age at emigration; middle) of emigration, and relative timing of emigration by age (right) for fish in tributaries of Grimwith (top) and Langsett (bottom) reservoirs.

### 3.2 Reservoir movements

Home range (K95 ± S.D.) was largest during October (24.9 ± 8.29 ha), November (26.9 ± 6.69 ha) and December (26.9 ± 3.6 ha) and smallest in February (7.7 ± 2.86 ha) and March (7.0 ± 3.45 ha) (Figure 3). There was a highly significant difference in mean brown trout activity between months (Scheirer-Ray-Hare test: *X*^2^ = 39.99, *P* <0.01) and time of day (Scheirer-Ray-Hare test: *X*^2^ = 9.89, *P* =<0.01), peaking in May and at twilight, although the interaction between month and time of day was not significant (Scheirer-Ray-Hare test: *X*^2^ = 5.56, *P* >0.05) (Figure 4). Brown trout utilised a range of water depths, from the surface to a maximum of 27.7 m, with the majority (99%) in the upper 10 m (Figure 4). There was no difference in mean brown trout depth between months (Scheirer-Ray-Hare test: *X*^2^ = 1.25, *P* >0.05), but time of day was significant (Scheirer-Ray-Hare test: *X*^2^ = 21.24, *P* =<0.01), with fish moving between deep water during the day and shallower water at night, although the interaction between month and time of day was not significant (Scheirer-Ray-Hare test: *X*^2^ = 5.56, *P* >0.05) (Figure 4). All seven acoustic-tagged brown trout were detected approaching the weirs, but not further upstream, with five approaching both weirs, one approaching only the River Little Don and one only Thickwoods Brook.

**Figure 3.**
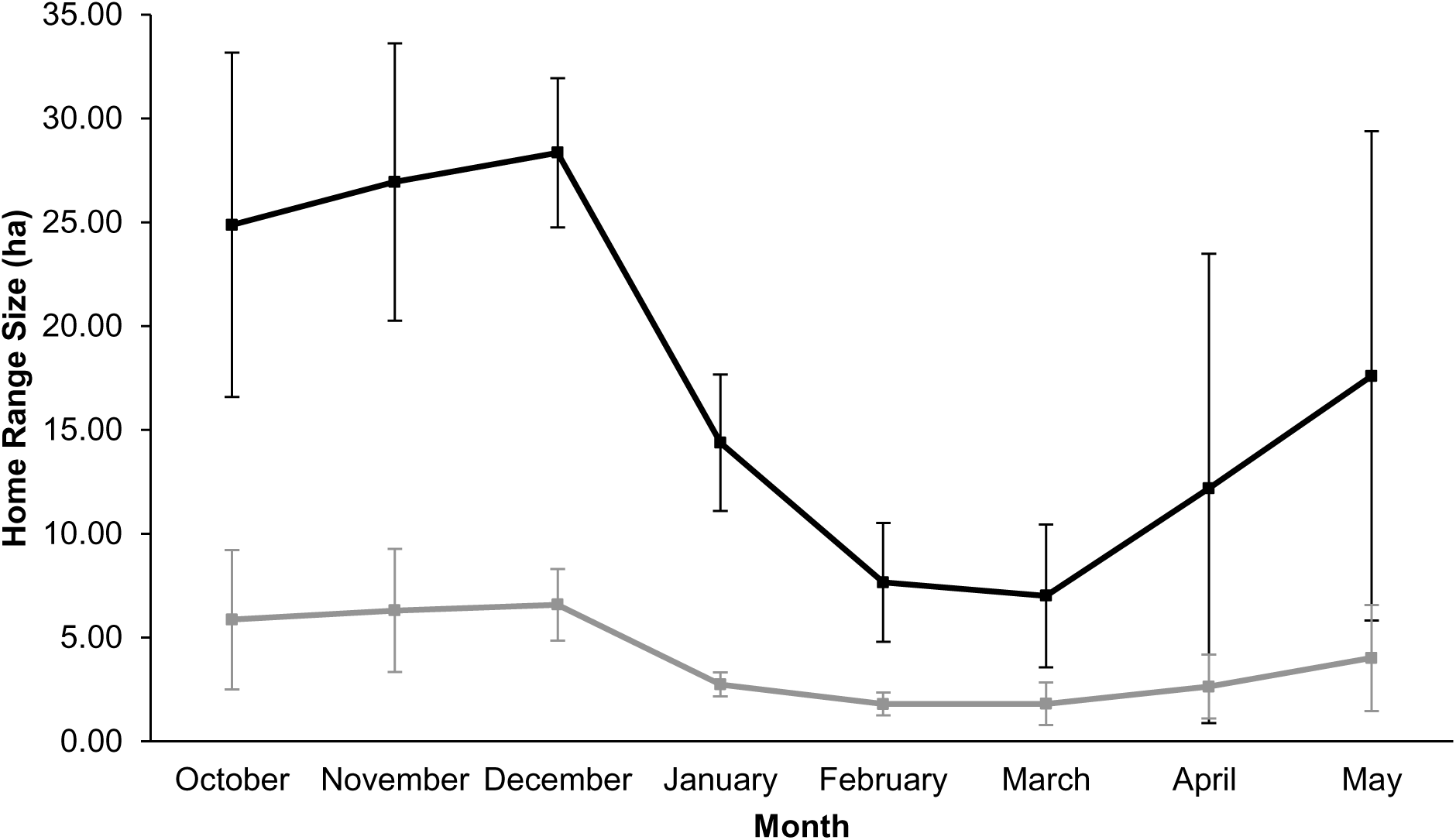
Mean (± S.D.) monthly K95 (black line) and K50 (grey line) home range size (hectares) of acoustic-tagged brown trout in Langsett Reservoir, October 2016–May 2017.

**Figure 4.**
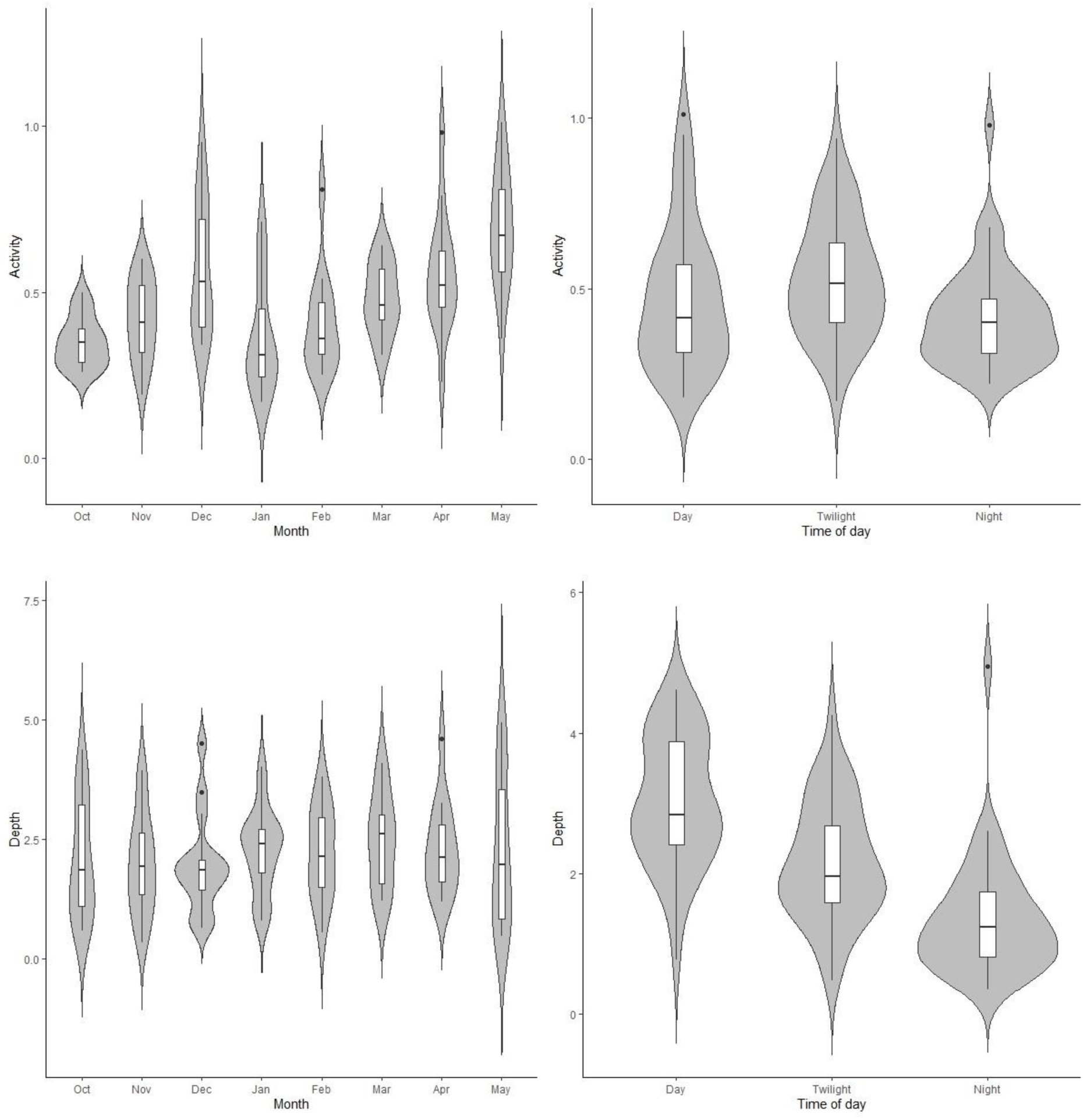
Violin plots of activity (m s^-2^; top) and depth (m; bottom) for each month (left) and time of day (right). Grey shaded areas represent the density of data for that value.

### 3.3 Upstream migration; tributary immigration

Across all years, 42.9% of fish tagged in Grimwith Reservoir and 37.2% of fish that emigrated from a tributary were detected moving upstream >30-days post-emigration (Figure 5). By contrast, 64.1% of fish tagged in Langsett Reservoir and 20.5% and 39.3% of fish that emigrated from River Little Don and Thickwoods Brook, respectively, were detected moving upstream (downstream of the weirs) into either one or both tributaries >30-days post-emigration (Figure 5). Fish tagged in reservoirs were not evenly distributed across tributaries during their upstream migration; the largest proportion entered Gate Up Gill (60%) at Grimwith Reservoir (Blea Gill Beck = 40%) and River Little Don (52.2%) at Lansgett Reservoir (Thickwoods Brook = 16.4% and 31.4% entered both). Of the fish that emigrated from each tributary, the proportion that subsequently returned or entered a different tributary was comparable at both reservoirs. For example, 44.4% of fish that emigrated from River Little Don returned, 44.4% entered Thickwoods Brook and 11.1% entered both. Ultimately, the majority of fish that entered River Little Don (92.0%; *n* = 173/188) and Thickwoods Brook (89.2%; *n* = 99/111) were tagged in Langsett Reservoir. Only three of 88 (3%) brown trout that approached the weir on River Little Don passed upstream prior to fish pass construction and three of 123 (2%) that approached the weir on Thickwoods Brook (throughout the study) passed upstream.

**Figure 5.**
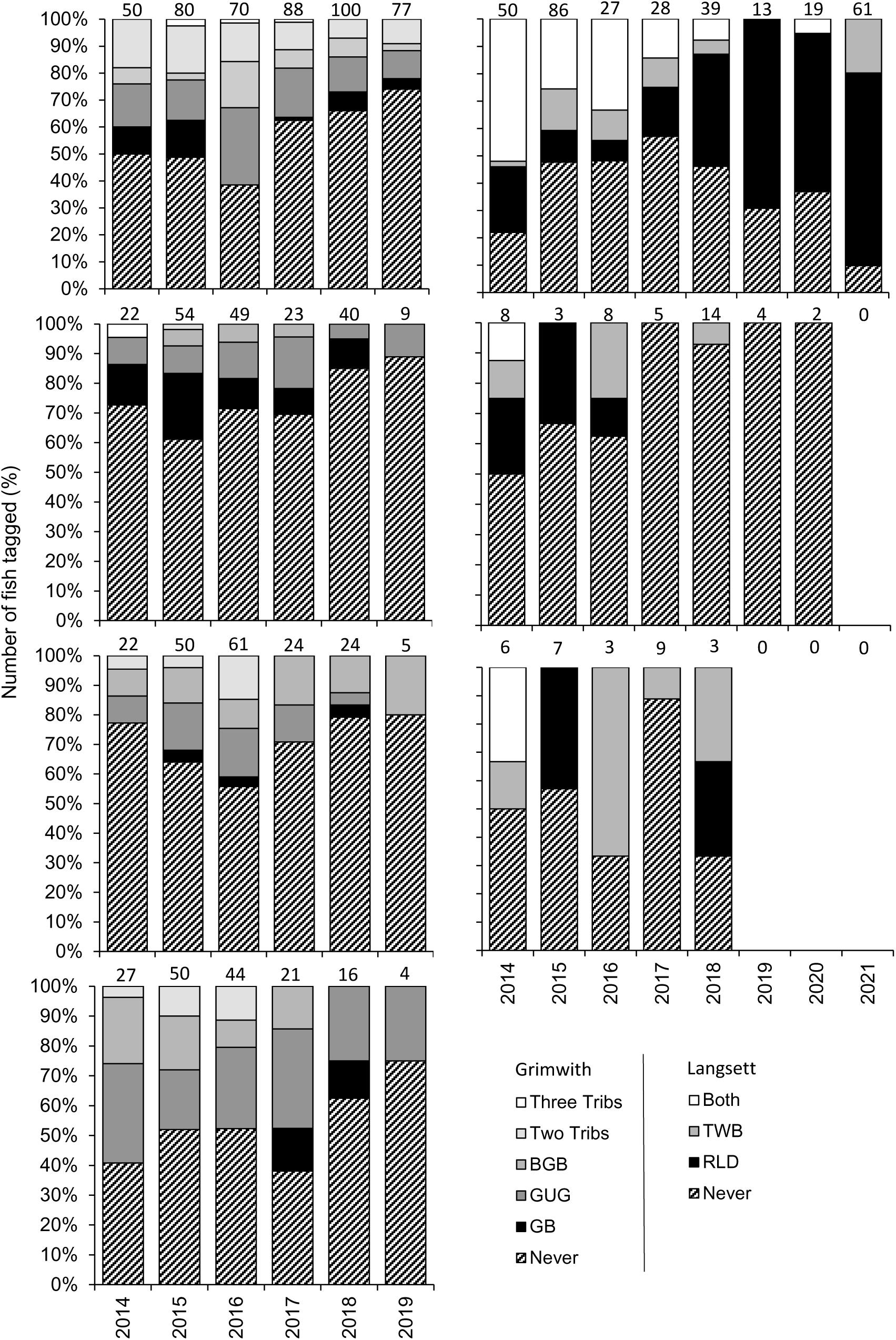
Proportion fish tagged (data label = total *n*) in each year at Grimwith (left) Reservoir (top left), Langsett (right) Reservoir (top right) or those that emigrated from Grimwith Beck (2^nd^ middle left), Gate Up Gill (3^rd^ middle left), Blea Gill Beck (bottom left), River Little Don (2^nd^ middle right) and Thickwoods Brook (3^rd^ middle right) detected immigrating into tributaries.

The modal age of fish entering tributaries was ≥4-year-old at Grimwith Reservoir and 1-year-old at Langsett Reservoir, influenced by age at tagging, with fish up to 7- and 10-years-old entering tributaries at the respective reservoirs (Figure 6). At Grimwith Reservoir, the median time between emigration from capture tributary to returning was 345.6 days (IQR=183.6 - 804.5, *n* = 120). Whereas at Langsett Reservoir, the median time between emigration from capture tributary to returning was 90.4 days (45.85 - 222.4, *n* = 16). At both reservoirs, peak immigration occurred in October-December, predominantly 3- and ≥4-year-old fish, although upstream movements occurred throughout the year and for all age classes (Figure 6).

**Figure 6.**
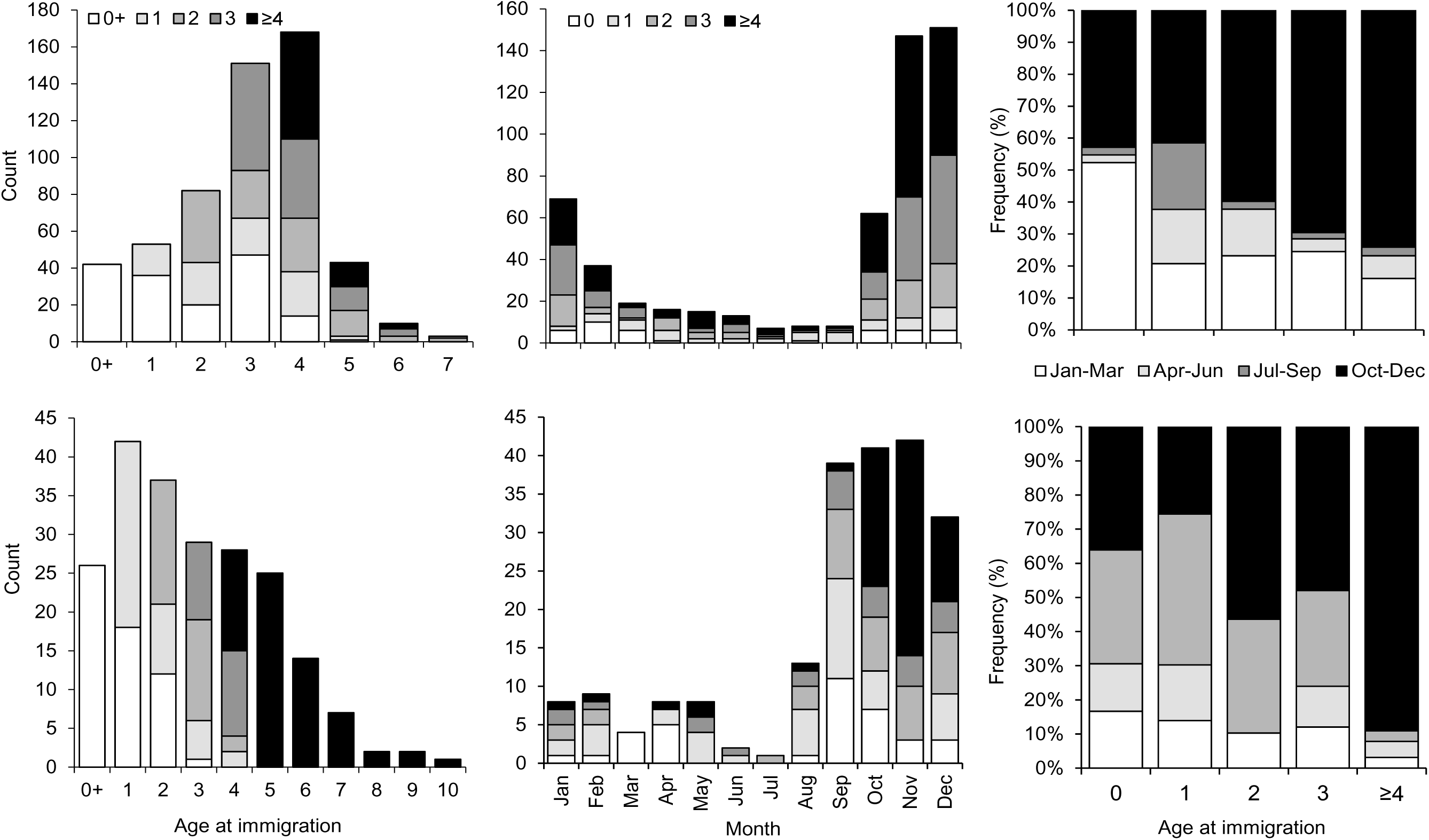
The age at immigration (left) and month of immigration (middle), including age at tagging, and relative timing of immigrations by age (right) for fish at Grimwith (top) and Langsett (bottom) reservoirs.

### 3.4 Fish pass efficiency

The total attraction, entrance, Larinier passage and FPS passage efficiencies for non-translocated fish approaching from a downstream direction were 30%, 69%, 36% and 14%, respectively, although there was interannual variability (Table 1). Fish that ascended the fish pass approached the weir a similar number times (median = 3: IQR = 1.75 – 4.25) to fish that did not pass (median = 2: IQR = 1 – 11.5) (Mann-Whitney *U* test: *Z* = -0.288, *n* = 115, *P* > 0.05). All non-translocated fish that approached the weir from a downstream direction and passed through the fish pass were caught and tagged in Langsett Reservoir; five and four fish that emigrated from River Little Don and Thickwoods Brook, respectively, approached the weir post-fish pass construction and were not detected ascending. By contrast, fish translocated from the River Little Don, upstream of the fish pass, had an overall passage efficiency of 43% (*n* = 9/21). A small proportion of 0-, 1- or ≥4-year-old non-translocated fish were detected at the entrance to the fish pass, whereas larger proportions of 2- and 3-year-old fish tended not to approach (FP1) and enter (FP2) the fish pass (Figure 7). The overall passage efficiency for 2 and 3-year-old non-translocated (32%) and translocated (33%) fish was comparable (Mann-Whitney U test: Z = 0.148, n = 65, P = 0.882) (Figure 7). The median (IQR) time to pass for non-translocated (1.69 (0.35-9.28) days) and translocated (1.04 (0.80-97.07) days) fish was also comparable (Mann-Whitney U test: Z = 0.494, n = 20, P = 0.656). Weir approach (A4: range = Q_100_-Q_0.5_), fish pass attraction (FP1: Q_100_-Q_27.5_), entrance (FP2: Q_100_-Q_28.5_) and passage (FP3: Q_100_-Q_6.2_) occurred at decreasing river levels for both non-translocated and translocated fish (Figure 8). In addition, 80% of the fish that entered the pass when depths were higher than Q_43.7_ did not pass. No fish tagged in the River Little Don were detected moving through the fish pass in a downstream direction

**Figure 7.**
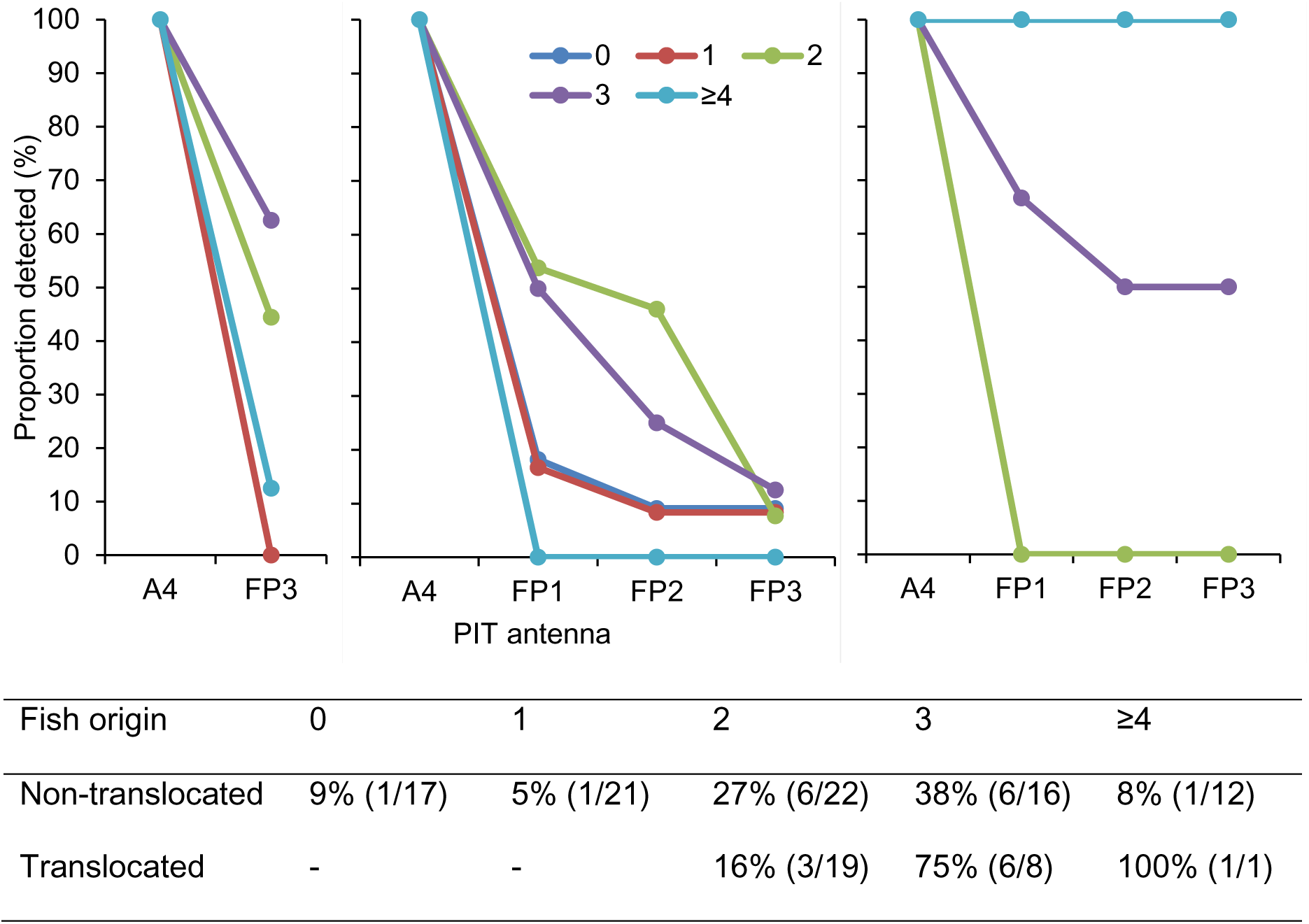
Percentage of each fish age class detected at approach (A4), attraction (FP1), entrance (FP2) and passage (FP3) PIT antennas for non-translocated fish in 2017-20 (left), non-translocated fish in 2020-22 (middle) and translocated fish in 2020-21 (right), and overall passage efficiency by age (bottom).

**Figure 8.**
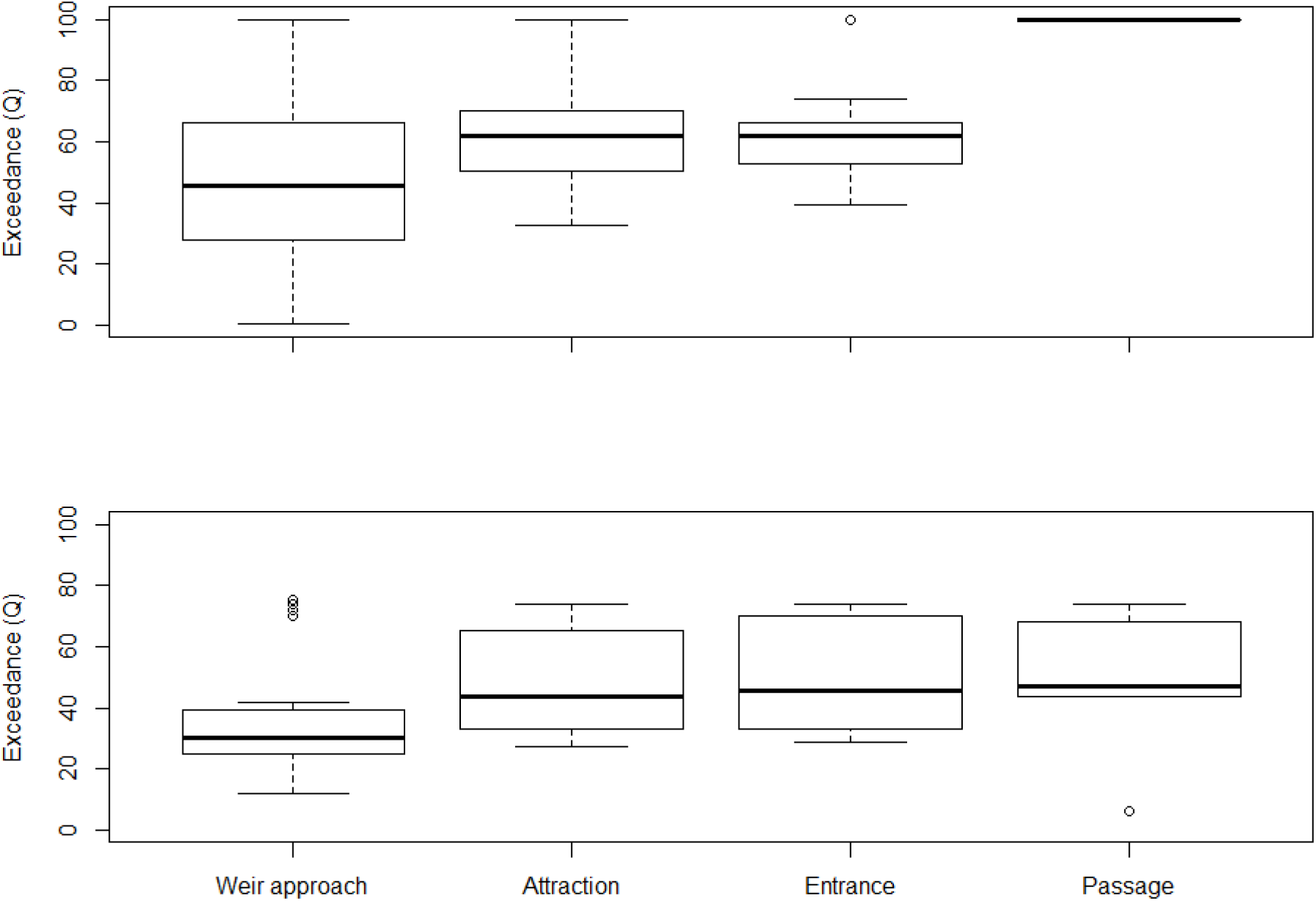
River depth exceedance (Q_x_) during weir approach (A4), fish pass attraction (FP1), entrance (FP2) and passage (FP3) for non-translocated (top) and translocated (bottom) fish in 2020/21.

**Table 1.** Annual and total fish passage efficiency (confidence intervals; *n*) and passage time metrics for non-translocated (left) and translocated (right) fish that approached from downstream. – denotes periods where antennas were not in-situ to allow calculations.

| Metric (antenna) | 2017/18 | 2018/19 | 2019/20 | 2020/21 | 2021/22 | Total (CI; *n*) | 2019/20 | 2020/21 | Total (CI; *n*) |
| --- | --- | --- | --- | --- | --- | --- | --- | --- | --- |
| Available fish (A4) | 16 | 15 | 13 | 14 | 39 | 97 | 19 | 9 | 28 |
| Attraction efficiency (FP1/A4) | - | - | - | 29%<br>(4/14) | 31%<br>(12/39) | 30% (20-44;<br>16/53) | - | 56%<br>(5/9) | 56% (26-81;<br>5/9) |
| Entrance efficiency (FP2/FP1) | - | - | - | 100%<br>(4/4) | 58%<br>(7/12) | 69% (44-86;<br>11/16) | - | 80%<br>(4/5) | 80% (36-96;<br>4/5) |
| Larinier passage efficiency (FP3/FP1) | - | - | - | 25%<br>(1/4) | 43%<br>(3/7) | 36% (15-65;<br>4/11) | - | 100%<br>(4/4) | 100% (48-99;<br>4/4) |
| FPS passage efficiency (FP3/A4) | 31%<br>(5/16) | 7%<br>(1/15) | 31%<br>(4/13) | 7%<br>(1/14) | 8%<br>(3/39) | 14% (9-23;<br>14/97) | 5%<br>(1/19) | 56%<br>(5/9) | 21% (10-40;<br>6/28) |
| Overall passage efficiency (FP3 or A3/A4) | 31%<br>(5/16) | 7%<br>(1/15) | 31%<br>(4/13) | 14%<br>(2/14) | 8%<br>(3/39) | 15% (10-24;<br>15/97) | 11%<br>(2/19) | 78%<br>(7/9) | 32% (18-51;<br>9/28) |

### 3.5 Population-scale impact of fish pass construction

Geometric mean densities of brown trout in the River Little Don (impact site) and control reaches both decreased after the fish pass opened (Figure 9). Geometric mean density of 0+ brown trout declined by 68% between the before and after period in the River Little Don and by 73% in the control sites. The decline in 0+ density in the impact site was significantly less than that in the control sites (Temporal difference (Z) = 0.772, S.E. 0.253, 95% C.I. = 0.529, *P*<0.05), i.e., the fish pass has had a positive effect on 0+ brown trout densities in the River Little Don. Geometric mean density of >0+ brown trout declined by 40% between the before and after period in the River Little Don and by 32% in the control sites. There was no significant difference in the declines between the before and after period in the >0+ brown trout mean density at the impact site relative to the control sites (Temporal difference (Z) = -0.007, S.E. = 0.1326, 95% C.I. = 0.2775, *P* = >0.05).

**Figure 9:**
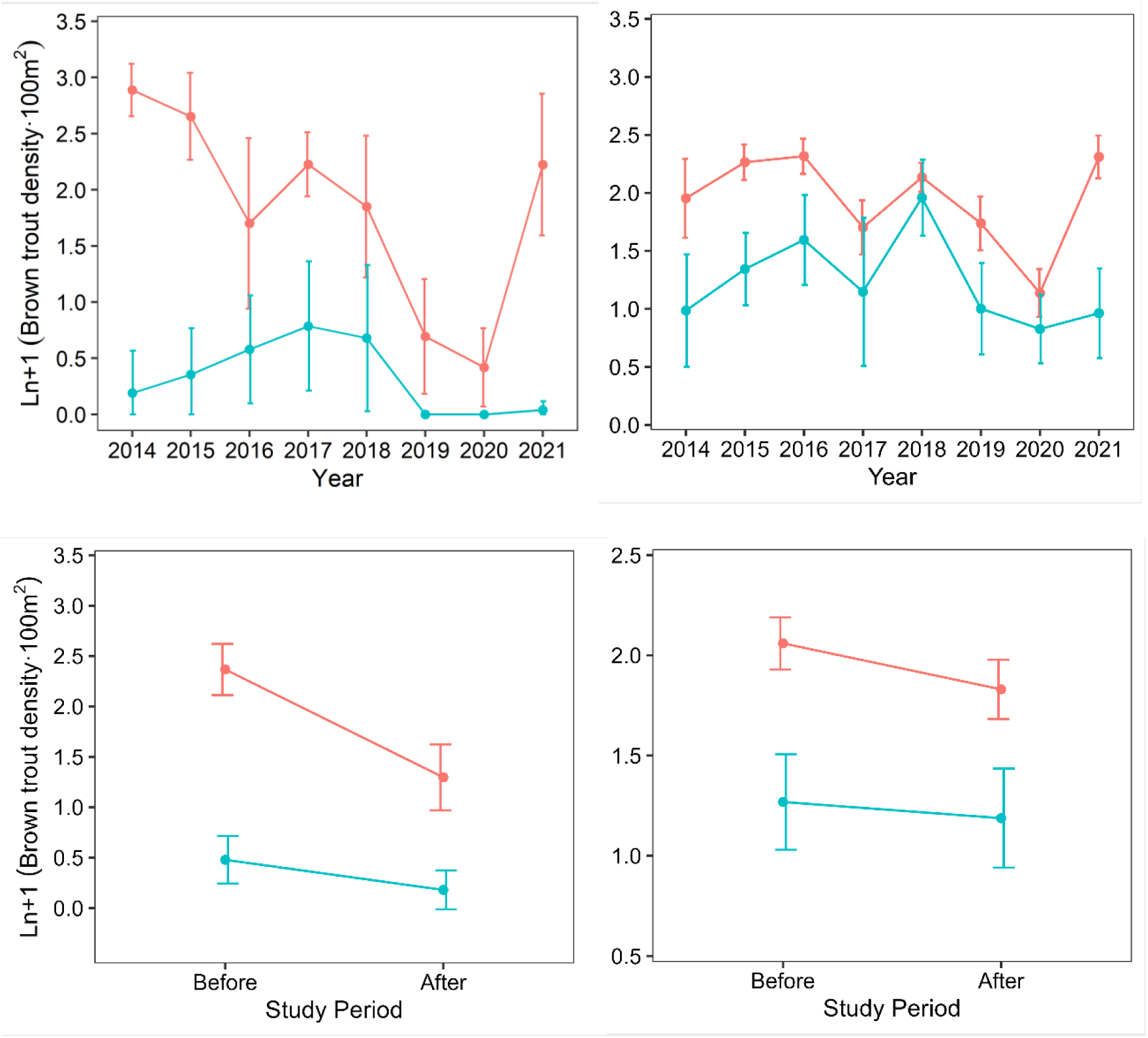
Mean (95% confidence intervals) annual (top) and before (2014-2017) versus after (2018-2021) fish pass opening (bottom) Ln+1 density of 0+ (left) and >0+ (after) brown trout for control (red) and impact sites (blue).

## 4. DISCUSSION

This investigation compared the spatial ecology and population dynamics of brown trout between reservoirs with (impact; Langsett Reservoir) and without (control; Grimwith Reservoir) barriers to fish movements into afferent headwater tributaries. Fish that entered Langsett Reservoir from afferent tributaries were unable to return and contribute to future populations due to the barriers. Connectivity was remediated (i.e., Larinier fish pass) for a moderate proportion of fish that approached a weir on one tributary (River Little Don), and although population densities in the after period were lower at both impact and control sites the BACI analysis indicated a significant positive effect of the fish pass on 0+ density in the River Little Don. Here we discuss how telemetry methods were key to providing a holistic understanding of brown trout ecology in headwater reservoirs, including downstream and upstream migrations in afferent tributaries, space use and activity in the reservoir, and anthropogenic influences on connectivity.

Brown trout in the tributaries of Grimwith and Langsett reservoirs predominantly emigrated at comparable ages, between 0-2 and 1-3 years of age, respectively, and up to the age of 5 and 6-years-old. Craig (1982) found that brown trout in the tributaries of Lake Windermere, UK, migrate downstream in their first to third years, and thereafter remain river-resident. While Jonsson *et al*. (1999) also found the majority of brown trout migrating downstream into Lake Femund, Norway, were two or three years old, although individuals as old as eight were recorded moving downstream in small number. Brown trout moved downstream in autumn, presumably to seek refuge in deeper water during winter, and in spring, presumably to access food resources in the summer, similar to the findings of Stuart (1953), Lien (1979) and Jonsson & Jonsson (2011). It was concluded that brown trout performed active downstream migration rather than displacement, passive drift or washout, given the weirs at Langsett Reservoir (Impact; emigration rate = 26%) appeared to thwart emigration relative to Grimwith Reservoir (Control; emigration rate = 85%). Although, brown trout populations in Langsett Reservoir tributaries were low, and Olsson & Greenberg (2004) found lower than expected population levels dissuaded resident brown trout from migrating downstream due to plentiful food and habitat availability.

Adult brown trout in the reservoir had larger home ranges during the spawning period (October-December) than any other month (January-May). Ovidio *et al*. (2002) also reported the largest home range of river-resident brown trout was during the spawning migration. Winter reductions in home range size of lake trout (*Salvelinus namaycush*) and lake-dwelling rainbow trout (*Oncorhynchus mykiss*) have been attributed to the cooler temperatures reducing metabolic rate of the fish, ambient light levels and day length (Blanchfield *et al*., 2009; Watson *et al*., 2019). Here, brown trout were most active in May, probably associated with feeding, due to increases in both food availability and warmer temperatures (Ojanguren *et al*., 2001; Mierzejewska et al., 2022). Throughout the study, trout were lower in the water column during daylight hours and closer to the surface during the hours of darkness (Bardonnet *et al*., 2006; Nash *et al*., 2022), potentially associated with feeding (Dervo *et al*., 1991) or may coincide with offshore-onshore movements (Cote *et al*., 2020). All seven acoustic tagged brown trout approached the man-made weirs in the headwater tributaries at Langsett Reservoir, with five moving between the two tributaries, predominantly during November and December presumably in search of suitable spawning habitat.

Brown trout predominantly moved from Grimwith (Control) and Lansgett (Impact) reservoirs into headwater tributaries between October and January, and tended to be older fish, and thus was probably associated with an upstream spawning migration (Piecuch *et al*., 2007; Jonsson & Jonsson, 2011). Upstream movements into the headwater tributaries occurred throughout the year, including for immature trout, and so spawning was not the sole driver, as also found by Carlsson *et al*. (2004). Crucially, only three upstream migrating fish were detected upstream of the weir on the River Little Don prior to fish pass construction with similar numbers able to ascend the weir on Thickwoods Brook, both headwater tributaries at Langsett Reservoir. Such disruption to spawning migrations can cause loss of fitness due to repeated attempts to pass impoundments, or increased time spent in sub-optimal conditions where the bottleneck occurs (Aarestrup & Jensen, 1998; Gerlier & Roche, 1998). They may also have to settle for less favourable spawning habitat or areas that become inundated when the reservoir fills over winter, which may lead to higher egg mortality (Battin, 2004; Thaulow *et al*., 2014). Alternatively, brown trout may have spawned in the reservoir, as found when there is a groundwater influx (Thaulow *et al*., 2014; Arostegui & Quinn, 2019; Brabrand *et al*., 2002), but we found no evidence of that here. Ultimately, fragmentation of habitats, independent of habitat loss, can increase the probability of extinction in populations (Fahrig, 2003).

The FPS passage efficiency for non-translocated brown trout was low (14%) but comparable to fish pass performance data for river-resident brown trout elsewhere (Dodd et al 2018; Lothian *et al.,* 2020; Bravo-Córdoba *et al*., 2022). It was largely attributed to fish that approached the weir not being detected at the entrance (attraction efficiency = 30%) and in turn not swimming up the pass (Larinier passage efficiency = 36%). The former may be attributed to the pre-barrage (Dodd *et al.,* 2023) or Larinier entrance extending beyond the weir face and the latter may be to the narrow width, gradient or length (Bunt *et al*., 2012; Noonan *et al*., 2012).Fish age and prevailing river level also influenced the passage rate. The passage rates for both non-translocated and translocated fish were highest at 2 and 3 years old, which corresponds to mature fish migrating into the River Little Don to spawn (Jonsson and Jonsson, 2011). Translocated fish may also have been homing to natal spawning grounds (Gosset et al. 2006). Conversely, 0- and 1-year old fish detected approaching the weir may have been merely living in the reach and were not motivated to pass (Dodd *et al*., 2023). While ≥4-year-old fish may have had fidelity to a previous spawning location in the short reach of river downstream of the weir (∼50 m when the reservoir is full) prior to fish pass opening (Pess *et al*., 2014; Davies *et al*., 2023). All fish pass approaches were at depths <Q_27_, and thus elevated river levels may have caused turbulent flows in the vicinity of the weir which prevented the fish from locating the fish pass entrance (Bunt *et al*., 2000). Likewise, 80% of the fish that entered the pass when depths were higher than Q_43.7_ did not pass, perhaps because flows exceeded swimming capabilities (Bunt *et al*., 2012; Noonan *et al*., 2012). There were also no instances of fish emigrating from River Little Don moving through the pass in a downstream direction, despite the importance of bidirectional connectivity (Calles & Greenberg, 2009; Bravo-Córdoba et al., 2023).

Unlike Grimwith Reservoir (control), the brown trout populations in the headwater tributaries at Langsett Reservoir (impact) were upstream of largely insurmountable barriers (pre-fish pass construction) and must be sustained by brown trout that do not move downstream over the weirs. It was hence speculated that fragmentation could lead to population declines, as reported by Gosset *et al*., (2006). Fish pass construction did not culminate in an increase in 0+ or >0+ brown trout in the River Little Don. However, the decline of 0+ mean density of brown trout at the impact site was significantly less than that of the control site. Other studies have reported population recovery following efforts to remediate connectivity for brown trout (Birnie-Gauvin *et al*., 2018; Duda *et al.,* 2021; Sun *et al.,* 2022) and sea lamprey (Pereira *et al*. 2017). Wittum *et al*. (2023) reported shifts in fish assemblage structure following two dam removals on the Penobscot River, North America, but alosines were absent from the upper river due to the requirement to use fish passes at two dams. It may be that too few fish used the fish pass to culminate in a population scale increase in brown trout recruitment in the River Little Don. Alternatively, physical habitat, environmental variables (river level and temperature) and/or density dependent mortality had an overarching influence on brown trout recruitment (Lobón-Cerviá and Mortensen 2005; Lobón-Cerviá 2009). Notwithstanding, the genetic diversity upstream of the weir was impacted (Moccetti *et al*., 2023), and the investigation occurred over relatively short timeframe and thus population increases may occur more incrementally over a longer timeframe. Even occasional passages could be sufficient for maintaining a population or re-establishing one after an extinction event (Mahlum et al., 2018).

### Conclusions

This study combined PIT and acoustic telemetry to provide a long-term and holistic understanding of brown trout movement ecology in headwater reservoirs, and incorporated a before/after control/impact study design to quantify the impact of a fish pass construction. Emigration from and immigration into afferent tributaries was performed by all age classes of brown trout and occurred throughout the year, but both were impeded by man-made weirs at the impact reservoir. Fish in the reservoir had the largest home range in October – December, with activity influenced by both month and time of day, and fish occupied shallow depths (relative to reservoir depth), especially at night. Fish pass construction remediated connectivity was for certain age classes of fish (2- and 3-year-old), and although the BACI indicated a positive effect on 0+ trout density, the overall abundance of brown trout in the River Little Don did not increase in the four years following the opening of the fish pass. Overall, this investigation significantly furthers our understanding of brown trout spatial ecology and population dynamics in reservoirs and headwater tributaries.

## ACKNOWLEDGEMENTS

Many thanks to University of Hull International Fisheries Institute (HIFI) staff and post-graduate student for their assistance collecting field data. I would also like to thank Yorkshire Water Services for supporting the research and providing access to the study reservoirs and rivers.

## 5. CONTRIBUTIONS

Conceptualization: JDB, JPH Data curation: JRD, HO, LW Formal analysis: JRD, HO

Funding acquisition: JDB, JPH, JRD, BG Investigation: JRD, JDB, LW, ADN, RAAN, JPH, PM Methodology: JDB, JRD

Supervision: JDB, RAAN Writing – original draft: JRD

Writing – review & editing: JRD, JDB, ADN, RAAN, PM, BG, DAJ

## 6. SUPPLEMENTARY MATERIAL

**Table 3:**
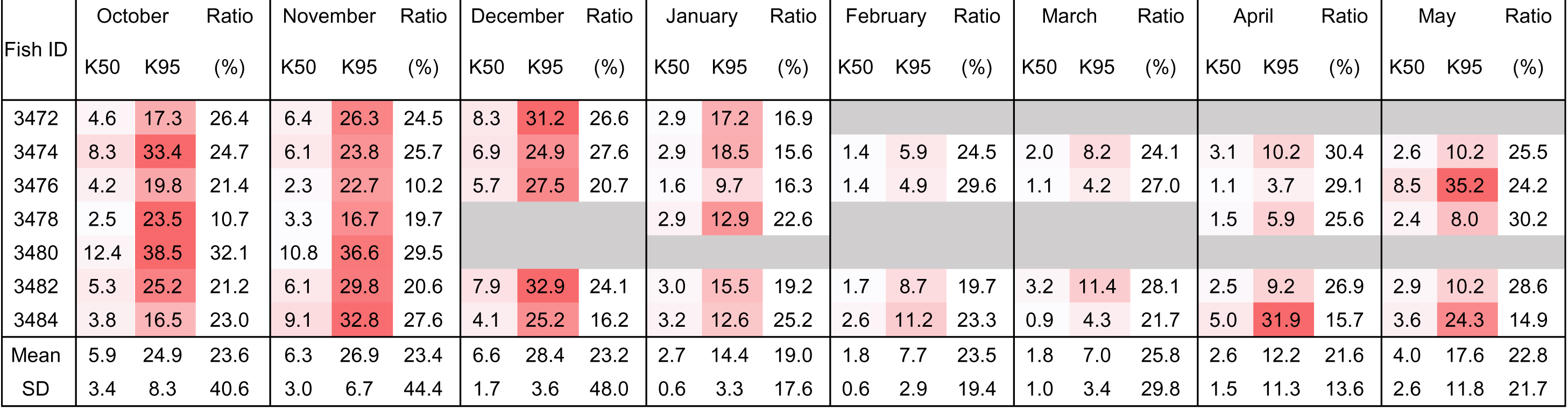
Fish ID, monthly K50 and K95 home range size (hectares) and K50/ K95 ratio (%) of fish in Langsett reservoir October 2016 - May 2017 and mean home ranges. Conditional formatting highlights the months with the larger home range sizes.

**Table 4:** Activity levels (ms^-2^) of tagged brown trout in Langsett reservoir during day, twilight and night. Shading indicates high levels of activity for each fish.

| Fish<br>Number | October |  |  | November |  |  | December |  |  | January |  |  | February |  |  | March |  |  | April |  |  | May |  |  |
| --- | --- | --- | --- | --- | --- | --- | --- | --- | --- | --- | --- | --- | --- | --- | --- | --- | --- | --- | --- | --- | --- | --- | --- | --- |
|  | D | T | N | D | T | N | D | T | N | D | T | N | D | T | N | D | T | N | D | T | N | D | T | N |
| 3472 | 0.26 | 0.32 | 0.29 | 0.57 | 0.50 | 0.31 | 0.95 | 0.81 | 0.54 |  |  |  |  |  |  |  |  |  |  |  |  |  |  |  |
| 3474 | 0.29 | 0.36 | 0.31 | 0.19 | 0.33 | 0.31 | 0.39 | 0.40 | 0.37 | 0.31 | 0.33 | 0.22 | 0.47 | 0.54 | 0.32 | 0.57 | 0.64 | 0.41 | 0.50 | 0.62 | 0.44 | 0.81 | 0.78 | 0.55 |
| 3476 | 0.50 | 0.46 | 0.39 | 0.59 | 0.56 | 0.46 | 0.84 | 0.71 | 0.53 | 0.66 | 0.71 | 0.39 | 0.37 | 0.81 | 0.47 | 0.42 | 0.45 | 0.31 | 0.40 | 0.79 | 0.50 | 1.01 | 0.94 |  |
| 3478 | 0.27 | 0.26 | 0.28 | 0.26 | 0.42 | 0.41 |  |  |  | 0.18 | 0.17 | 0.31 |  |  |  |  |  |  | 0.23 | 0.68 | 0.98 | 0.53 | 0.52 | 0.67 |
| 3480 | 0.36 | 0.35 | 0.26 | 0.52 | 0.52 | 0.41 |  |  |  |  |  |  |  |  |  |  |  |  |  |  |  |  |  |  |
| 3482 | 0.32 | 0.47 | 0.45 | 0.49 | 0.60 | 0.41 | 0.73 | 0.71 | 0.45 | 0.57 | 0.51 | 0.29 | 0.32 | 0.45 | 0.25 | 0.53 | 0.62 | 0.46 | 0.63 | 0.55 | 0.52 | 0.67 | 0.57 | 0.36 |
| 3484 | 0.38 | 0.40 | 0.36 | 0.24 | 0.40 | 0.32 | 0.39 | 0.46 | 0.34 | 0.26 | 0.32 | 0.23 | 0.29 | 0.35 | 0.26 | 0.34 | 0.46 | 0.57 | 0.41 | 0.61 | 0.47 | 0.87 | 0.81 | 0.68 |
| **Mean** | 0.34 | 0.37 | 0.33 | 0.41 | 0.48 | 0.37 | 0.66 | 0.62 | 0.44 | 0.39 | 0.41 | 0.29 | 0.36 | 0.54 | 0.33 | 0.46 | 0.54 | 0.44 | 0.43 | 0.65 | 0.58 | 0.78 | 0.73 | 0.57 |
| **STDev** | 0.08 | 0.07 | 0.07 | 0.17 | 0.10 | 0.06 | 0.26 | 0.18 | 0.09 | 0.21 | 0.21 | 0.07 | 0.08 | 0.20 | 0.10 | 0.11 | 0.10 | 0.11 | 0.15 | 0.09 | 0.22 | 0.18 | 0.18 | 0.15 |

**Table 5:** Mean monthly depth (m) during day (D), twilight (T) and night (N) for brown trout in Langsett reservoir, October 2016 to May 2017. With conditional formatting highlighting the greatest depths for each month (colour scale; dark red = greatest depth, no shading = shallowest depth, grey = no data).

| Fish Number | October |  |  | November |  |  | December |  |  | January |  |  | February |  |  | March |  |  | April |  |  | May |  |  |
| --- | --- | --- | --- | --- | --- | --- | --- | --- | --- | --- | --- | --- | --- | --- | --- | --- | --- | --- | --- | --- | --- | --- | --- | --- |
|  | D | T | N | D | T | N | D | T | N | D | T | N | D | T | N | D | T | N | D | T | N | D | T | N |
| 3472 | 2.71 | 1.88 | 1.26 | 3.05 | 2.27 | 0.86 | 2.06 | 1.69 | 0.77 |  |  |  |  |  |  |  |  |  |  |  |  |  |  |  |
| 3474 | 3.98 | 3.22 | 1.33 | 3.89 | 2.70 | 1.35 | 4.50 | 3.49 | 1.91 | 4.02 | 3.52 | 2.53 | 3.80 | 3.01 | 1.77 | 4.09 | 2.90 | 1.64 | 4.61 | 3.20 | 2.06 | 4.49 | 2.93 | 1.79 |
| 3476 | 1.55 | 0.89 | 0.67 | 2.08 | 1.37 | 0.75 | 2.09 | 1.63 | 0.88 | 2.41 | 1.73 | 0.80 | 2.94 | 2.36 | 1.43 | 3.33 | 2.20 | 1.21 | 3.26 | 2.24 | 2.19 | 2.65 | 1.97 |  |
| 3478 | 1.43 | 1.10 | 0.60 | 2.23 | 1.61 | 0.35 |  |  |  | 2.68 | 2.65 | 1.10 |  |  |  |  |  |  | 2.70 | 1.79 |  | 2.27 | 0.67 | 0.83 |
| 3480 | 3.32 | 2.07 | 0.63 | 3.94 | 2.64 | 1.51 |  |  |  |  |  |  |  |  |  |  |  |  |  |  |  |  |  |  |
| 3482 | 4.38 | 3.49 | 2.45 | 3.56 | 1.94 | 1.04 | 3.02 | 1.86 | 1.31 | 2.75 | 2.41 | 1.88 | 3.30 | 1.93 | 1.22 | 3.81 | 2.61 | 2.60 | 2.03 | 1.20 | 1.26 | 0.77 | 0.47 | 0.54 |
| 3484 | 3.88 | 3.35 | 1.85 | 2.52 | 1.51 | 0.48 | 2.03 | 1.57 | 0.65 | 2.76 | 1.92 | 0.98 | 2.77 | 1.51 | 0.56 | 2.63 | 1.33 | 1.25 | 2.83 | 1.55 | 1.31 | 4.17 | 4.25 | 4.95 |
| **Mean** | 3.03 | 2.29 | 1.26 | 3.04 | 2.01 | 0.91 | 2.74 | 2.05 | 1.10 | 2.92 | 2.45 | 1.46 | 3.20 | 2.21 | 1.25 | 3.47 | 2.26 | 1.68 | 3.09 | 1.99 | 1.70 | 2.87 | 2.06 | 2.03 |
| **STDev** | 1.18 | 1.08 | 0.70 | 0.78 | 0.54 | 0.43 | 1.07 | 0.81 | 0.52 | 0.63 | 0.71 | 0.73 | 0.46 | 0.64 | 0.51 | 0.64 | 0.68 | 0.65 | 0.96 | 0.77 | 0.49 | 1.51 | 1.58 | 2.02 |

**Figure 6:**
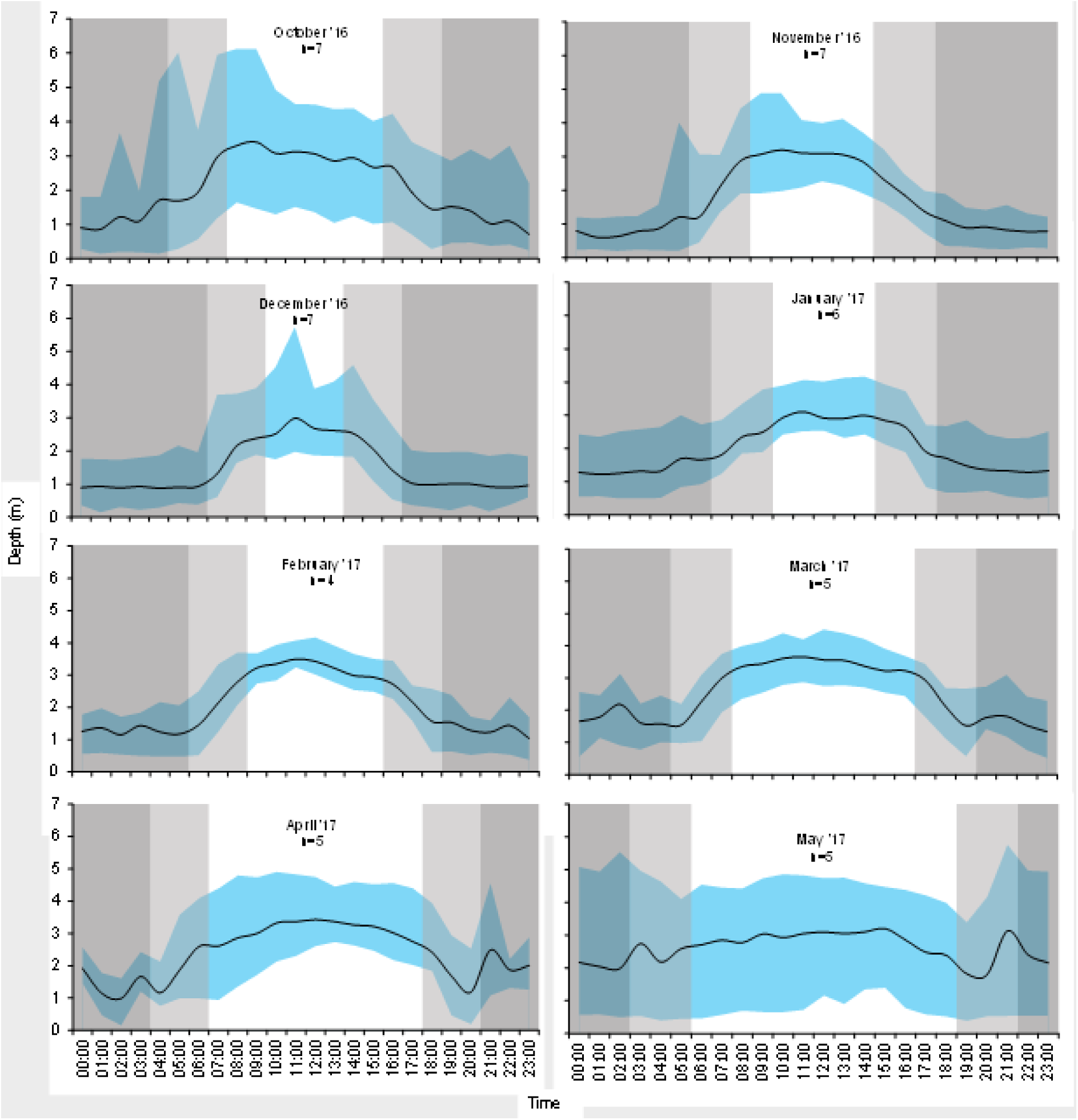
Mean depths (m) of tagged brown trout in Langsett reservoir. Minimum and maximum mean depths illustrated by blue shaded region, dark grey = night, light grey= twilight, no shading = day (n=number of fish).

**Table 2:** 0+ and ≥0+ brown trout density (fish per 100 m^2^) and abundance classification for Langsett and Grimwith reservoirs in 2014 – 2021.

| River Name | Site | 0+ brown trout |  |  |  |  |  |  |  | ≥0+ brown trout |  |  |  |  |  |  |  |
| --- | --- | --- | --- | --- | --- | --- | --- | --- | --- | --- | --- | --- | --- | --- | --- | --- | --- |
|  | identifier | 2014 | 2015 | 2016 | 2017 | 2018 | 2019 | 2020 | 2021 | 2014 | 2015 | 2016 | 2017 | 2018 | 2019 | 2020 | 2021 |
| Little Don | RLD1 | 0.00 | 0.00 | 0.00 | 0.00 | 0.00 | 0.00 | 0.00 | 0 | 1.30 | 1.51 | 1.27 | 0.00 | 5.00 | 0.50 | 1.42 | 1.05 |
|  | RLD2 | 0.00 | 0.00 | 0.00 | 0.00 | 0.98 | 0.00 | 0.00 | 0 | 1.07 | 2.04 | 2.96 | 3.80 | 5.56 | 2.41 | 1.33 | 0.60 |
|  | RLD3 | 0.00 | 0.00 | 0.76 | 1.52 | 7.36 | 0.00 | 0.00 | 0 | 0.40 | 1.77 | 4.55 | 1.52 | 4.33 | 0.44 | 0.27 | 1.76 |
|  | RLD4 | 0.00 | 0.53 | 0.35 | 1.04 | 0.29 | 0.00 | 0.00 | 0 | 1.02 | 5.78 | 3.83 | 0.70 | 4.99 | 2.41 | 1.04 | 0.78 |
|  | RLD5 | 0.00 | 2.74 | 2.44 | 2.44 | 1.75 | 0.00 | 0.00 | 0.27 | 2.49 | 4.11 | 8.78 | 6.83 | 5.25 | 2.51 | 1.44 | 4.80 |
|  | RLD6 | 2.15 | 0.48 | 2.94 | 5.34 | 0.00 | 0.00 | 0.00 | 6.75 | 6.90 | 3.33 | 5.08 | 5.08 | 15.20 | 3.63 | 3.00 | 2.45 |
| Thickwoods Brook | TWB1 | 31.50 | 4.81 | 2.87 | 11.48 | 0.00 | 0.00 | 0.00 | 11.31 | 8.77 | 11.22 | 10.04 | 2.87 | 8.62 | 1.82 | 2.88 | 9.00 |
|  | TWB2 | 8.60 | 4.49 | 0.00 | 29.76 | 1.64 | 0.00 | 0.00 | 15.21 | 0.86 | 5.99 | 9.47 | 1.35 | 9.84 | 13.11 | 1.09 | 7.54 |
|  | TWB3 | 23.50 | 8.80 | 0.00 | 17.86 | 3.60 | 0.00 | 0.00 | 1.83 | 1.96 | 4.40 | 7.65 | 3.83 | 7.21 | 2.73 | 2.38 | 6.08 |
|  | TWB4 | 25.00 | 7.81 | 0.00 | 8.96 | 1.49 | 0.00 | 0.00 | 1.35 | 9.09 | 12.50 | 19.70 | 7.17 | 11.94 | 2.94 | 1.23 | 3.66 |
|  | TWB5 | 18.20 | 5.47 | 0.00 | 3.19 | 0.00 | 0.00 | 0.00 | 0 | 4.55 | 10.94 | 7.43 | 4.25 | 8.11 | 2.78 | 2.29 | 13.52 |
|  | TWB6 | 10.80 | 11.71 | 0.00 | 6.41 | 0.00 | 0.00 | 0.00 | 4.78 | 4.51 | 10.81 | 5.70 | 4.99 | 4.60 | 2.27 | 1.62 | 4.74 |
| Blea Gill Beck | BGB1 | 9.75 | 3.52 | 2.42 | 1.81 | 7.25 | 0.00 | 0.35 | 6.39 | 3.30 | 7.54 | 9.07 | 5.44 | 9.07 | 9.63 | 1.06 | 10.15 |
|  | BGB2 | 18.18 | 16.16 | 19.40 | 6.47 | 8.62 | 0.00 | 0.53 | 7.88 | 3.25 | 5.05 | 7.01 | 4.31 | 5.93 | 7.94 | 4.21 | 11.18 |
|  | BGB3 | 21.43 | 17.01 | 17.54 | 9.80 | 5.16 | 0.56 | 1.02 | 1.59 | 2.60 | 9.28 | 7.22 | 2.06 | 6.71 | 4.52 | 1.53 | 15.75 |
| Gate Up Gill | GUG1 | 6.21 | 18.97 | 2.18 | 6.54 | 10.89 | 0.00 | 0.00 | 5.86 | 12.41 | 5.80 | 7.99 | 2.90 | 4.36 | 3.80 | 0.50 | 14.32 |
|  | GUG2 | 10.43 | 31.55 | 10.03 | 7.80 | 13.37 | 3.43 | 0.00 | 19.95 | 13.39 | 8.33 | 6.68 | 3.90 | 7.80 | 5.71 | 1.92 | 10.88 |
|  | GUG3 | 24.17 | 14.46 | 8.84 | 6.31 | 12.62 | 2.83 | 0.00 | 58.36 | 14.50 | 12.72 | 10.10 | 3.79 | 10.52 | 6.13 | 1.91 | 8.48 |
| Grimwith Beck | GB1 | 22.03 | 54.76 | 52.75 | 11.72 | 29.30 | 6.90 | 1.65 | 57.71 | 7.34 | 11.53 | 16.61 | 12.70 | 7.81 | 6.90 | 6.04 | 11.48 |
|  | GB2 | 28.46 | 33.77 | 38.25 | 10.26 | 30.78 | 15.38 | 5.46 | 59.77 | 13.01 | 10.39 | 10.26 | 10.26 | 6.53 | 5.13 | 3.08 | 11.26 |
|  | GB3 | 20.69 | 30.19 | 31.81 | 7.95 | 32.61 | 8.67 | 6.45 |  | 13.00 | 8.84 | 9.54 | 6.36 | 6.36 | 4.00 | 3.87 | 7.03 |

**Table 3.** Site and number of PIT tagged brown trout in tributaries and reservoirs per year.

|  | **2014** | **2015** | **2016** | **2017** | **2018** | **2019** | **2020** | **2021** |
| --- | --- | --- | --- | --- | --- | --- | --- | --- |
| River Little Don | 50 | 100 | 100 | 100 | 101 | 66 | 22 | 27 |
| Thickwoods Brook | 50 | 100 | 31 | 83 | 33 | 13 | 9 | 54 |
| Langsett Reservoir | 50 | 86 | 27 | 28 | 39 | 13 | 19 | 61 |
| River Little Don<br>Displaced |  |  |  |  |  | 26 |  |  |
| Grimwith Beck | 50 | 100 | 100 | 54 | 100 | 61 | - | - |
| Gate Up Gill | 50 | 99 | 100 | 50 | 101 | 38 | - | - |
| Blea Gill Beck | 51 | 100 | 100 | 46 | 68 | 28 | - | - |
| Grimwith Reservoir | 50 | 80 | 70 | 88 | 100 | 77 | - | - |

## Notes

### Competing Interest Statement

The authors have declared no competing interest.

